# Ecological lifestyle and gill slit height across sharks

**DOI:** 10.1101/2024.01.17.574433

**Authors:** Wade J. Vander Wright, Jennifer S. Bigman, Anthony S. Iliou, Nicholas K. Dulvy

## Abstract

Morphology that is linked to metabolic rate – metabolic morphology – provides broad comparative insights into the physiological performance and ecological function of species. However, some metabolic morphological traits, such as gill surface area, require costly and lethal sampling. Measurements from anatomically-accurate drawings, such as those in field guides, offer the opportunity to understand physiological and ecological relationships without the need for physical, lethal sampling. Here, we assess the relationship between the metabolic physiology and ecology of nearly all extant sharks. Specifically, we examine the relationship between gill slit height and each of the three traits that comprise ecological lifestyle: activity, maximum size, and depth. We find that gill slit heights are positively related to activity (measured by the aspect ratio of the caudal fin) and maximum size but negatively related to depth. We also show that gill slit height is best explained by the suite of ecological lifestyle traits rather than any single trait. These results suggest that more active, larger, and shallower species have higher metabolic demands and that these greater metabolic demands can be estimated from external morphological and ecological traits. Our work demonstrates that meaningful ecophysiological relationships can be revealed through measurable metabolic morphological traits from anatomically-accurate drawings.

## 1 Introduction

Metabolic rate is thought to set the pace of life, from governing organismal developmental rates and life history traits to ecosystem processes such as biomass cycling (Brown et al. 2004). This fundamental rate is typically measured indirectly by oxygen consumption over time in a controlled laboratory setting and is difficult to measure for large-bodied, free-swimming organisms such as sharks (Prinzing et al. 2021, Payne et al. 2015). For sharks (and other fishes), the surface area of the gills is tightly correlated with metabolic rate such that individuals and species with large gill surface areas have higher oxygen consumption rates (i.e., metabolic rates) (Scheuffele et al. 2021, Wegner et al. 2012, Bigman et al. 2021). However, there are few estimates of metabolic rates for sharks and rays and gill surface area can be laborious to measure, taking more than 25 hours per individual (Bigman et al. 2018; Lyons et al. 2019; VanderWright et al. 2020). Here, we seek to develop new trait measures that encompass metabolic ecological function, but can be measured consistently for almost all sharks and rays (Streit and Bellwood 2022).

Gill slits are external openings of the parabranchial cavity in sharks and rays (Elasmobranchii) where exhalent oxygen-depleted water exits the body (Wegner 2015). The structure of the gill slit arises from the extension of interbranchial septum, which supports the gill filaments and surface area where oxygen uptake occurs (Wegner 2011, 2015). Thus, there is a direct functional morphological connection between the height of the external gill slit length and the corresponding internal gill surface area: shark gill slit height is related to gill surface areas such that species with larger gill surface area have longer mean gill slit heights (VanderWright et al. 2020). Specifically, gill slit height is positively correlated with (i) total gill surface area across species and (ii) parabranchial gill surface area (within an individual’s gill chambers or parabranchia) both within and across species (VanderWright et al. 2020). Additionally, gill slit height measurements are generally equivalent whether measured on live specimens or anatomically-accurate illustrations and can be made in a matter of minutes (Vanderwright et al. 2020).

Similar to metabolic rate, gill morphology, including the gill slit height is also correlated with activity, habitat (depth), and maximum size of a species, collectively termed ‘ecological lifestyle’ (Gray 1954, Killen et al. 2016, Bigman et al. 2018). The activity level of a species can be measured directly through swimming speed or indirectly through the caudal fin morphology (Harding et al. 2021, Jacoby et al. 2015, Sambilay 1990). The caudal fin aspect ratio (CFAR) is the ratio of *h* is the height of the caudal fin and *s* is the area of the caudal fin, and is calculated as *A* = *h*^2^/*s* (Figure 1a) is highly positively correlated with swimming speed in both teleosts and sharks (Sambilay 1990, Iliou et al. *2023*). Species with greater CFAR values are more active and have higher oxygen demands (i.e., metabolic rate; Killen et al. 2016, Campos et al. 2018). Similarly, species that are pelagic with shallower average depths generally have faster swimming speeds, higher metabolic rates, and larger gill surface area compared to deeper dwelling benthic species (Bigman et al. 2018, Killen et al. 2016, Andrzejaczek et al. 2022). The maximum size of a species also has a predictable relationship with metabolic rate (even beyond what can be accounted for by measurement body mass, or the mass of the individual whose metabolic rate was measured), where larger-bodied species have higher metabolic rates (Brown et al. 2004, Wong et al. 2021).

**Figure 1.**
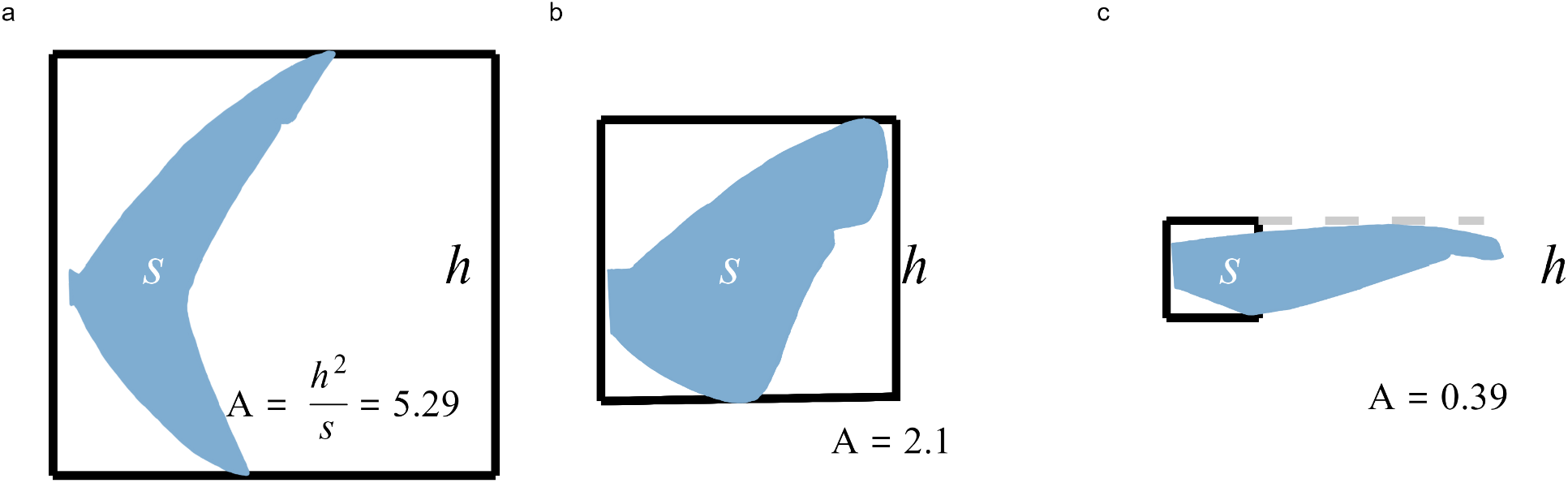
Schematic of how caudal fin aspect ratio (A) is calculated (*h*^*2*^*/s*). Black box represents the height of the caudal fin squared (*h*^*2*^) and blue tail shading are the surface areas (*s*). Panels show examples of the variation observed across shark species with high (Longfin Mako *Isurus paucus*; a) medium (Prickly Dogfish *Oxynotus bruniensis*; b), and low (Longfin Catshark *Apristurus herklotsi*; c) levels of activity.

Here, we examine whether gill slit height is broadly related to ecological lifestyle across sharks. Specifically we ask four questions: (1) do more active sharks (those with greater CFAR) have proportionately longer gill slits, (2) do larger-bodied sharks have proportionately longer gill slits, (3) do deeper-dwelling sharks have proportionately shorter gill slits, and finally (4) do all three ecological lifestyle traits together better explain variation in gill slit height than any individual component? As metabolic rate is related to ecological lifestyle and gill surface area is strongly correlated with metabolic rate, then GSH should be similarly related to ecological lifestyle. We expect that more active, larger-bodied, shallow water species will have greater GSH to accommodate greater oxygen demands (from higher metabolic rates). If true, rapid estimates of metabolic demand and by extension, life history traits, may be attainable without collecting live specimens or spending countless hours in laboratories.

## 2 Material and methods

### 2.1 Data collection

We measured total length (TL; snout to posterior of caudal fin), the height of each gill slit (following curvature), tail height (*h*; a vertical line from the top of the caudal fin to the bottom of the caudal fin), and tail area (*s*) from anatomically-accurate illustrations from *Sharks of the World*. This field guide is the most complete in terms of biodiversity and its illustrations have been validated for morphological accuracy (Ebert *et al*., 2013, Vanderwright et al. 2020, Bigman et al. 2018, Iliou et al. 2023). For each species of shark (*n* = 456), the lateral illustration was cropped and imported into ImageJ (cite for this) for measurement. Measurements of gill slit height were averaged across all gill slits (i.e., five, six, or seven gill slits) to generate a mean and divided by the TL of each species to calculate proportions for comparison (hereafter, ‘GSH’). We also calculated the summed height of all gill slits, to capture the full range of likely gill areas in six- and seven-gilled species. We omitted the Basking Shark (*Cetorhinus maximus*), Frilled sharks (Family Chlamydoselachidae), and the 20 Angel shark species (Family Squatinidae) as their gill slits were not completely visible in the illustrations. Caudal fin aspect ratio (CFAR) was calculated for each species defined as *A* = *h*^2^/*s*, where *h* is the height of the caudal fin and *s* is the area of the caudal fin (Palomares & Pauly 1989, Bigman et al. 2018; Figure 1).

Depth range (m) and maximum size (cm) were collated for each species using a combination of published IUCN Red List assessments (IUCN 2022), species checklists (Weigmann 2016), and *Sharks of the World* (Ebert *et al*. 2013).

### 2.2 Statistical analysis

To answer our questions, we fit four Bayesian linear models in a phylogenetic framework. Prior to analysis, GSH, CFAR, median depth, and maximum size were all natural-log transformed and all variables were standardized to facilitate model convergence and comparison.

Each ecological lifestyle trait was modeled as a predictor of GSH separately:

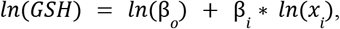

where GSH is the response variable -- the mean gill slit height of a species, β_o_ is the intercept, *x*_*i*_ is a given ecological lifestyle trait, and β_i_ is the slope of that ecological lifestyle trait. For Question 1 (do more active sharks have proportionately longer gill slits), *x*_*i*_ was CFAR, for Question 2 (do larger-bodied sharks have proportionately longer gill slits), *x*_*i*_ was maximum size, for Question 3 (do deeper-dwelling species have proportionately shorter gill slits), *x*_*i*_ was median depth, and for Question 4 (do all three ecological lifestyle traits together better explain gill slit height), the model included all three traits - CFAR, maximum size, and median depth (the ‘global model’). To be sure these related variables were not collinear in our model, we ensured the Variance Inflation Factor (VIF) was less than five (Quinn & Keogh 2002) (Table S4).

To account for the shared evolutionary history of traits between species, we included a random effect of phylogeny in all models. To do so, we pruned a taxon-complete chondrichthyan phylogenetic tree (a molecular tree of 615 species of sharks, rays, and chimaeras augmented with all remaining species based on taxonomic constraints; see Stein & Mull et al. (2018) for more detail) to the species in our dataset (n=456). To ensure our results were not biased due to using the infilled taxon-complete tree versus the molecular tree, we also fitted all models with a random effect of the molecular tree derived from the 615-species molecular tree trimmed to shark species with molecular data only (n=268).

A total of four models were fitted using the brms package in R v4.1.3 with four chains of 4000 iterations with 1000 warm-up iterations (5000 iterations total; Burkner 2017; R Core Team 2022). Uninformative priors were used and convergence was assessed by ensuring R-hat values (= 1 and effective sample size (ESS) >1000; Vehtari *et al*. 2019, Burkner 2017). All four models were compared to find the model with the most support and predictive ability using Pareto-smooth leave-one-out cross-validation (Vehtari et al. 2017). Models with larger expected log pointwise predictive densities (*elpd_loo*) values (and lower *looic* values) have more support and higher predictive ability (Vehtari et al. 2017). Generally, models with a difference of < 2 *looic* are considered to be equal in terms of fit and predictive ability and the most parsimonious model is usually preferred (Vehtari et al. 2017). Using the molecular versus full phylogenetic tree did not affect the model parameters substantially; however the lambda (*λ*) values were greater with the molecular tree (Table S1, S2).

## 3 Results

### Do more active sharks have larger gill slits?

More active sharks (those with greater caudal fin aspect ratios, CFAR) had larger mean gill slit heights (GSH; Figure 2). CFAR ranged from 0.25 in the Epaulette Shark (*Hemiscyllium ocellatum*) to 5.29 in the Longfin Mako (*Isurus paucus*) and their respective GSH values were 2.62% total length (TL) and 7.14% TL. An average shark species had a CFAR value of 1.05 and was estimated to have a GSH of 3% TL (Table 1). An increase in CFAR of one standard deviation (sd, i.e., 0.89 CFAR units) had an estimated increase in GSH of 0.37 sd (i.e., an increase of 0.44% TL) (Table 1).

**Table 1.** Model summary table comparing parameter estimates (with bayesian credible intervals), posterior standard deviation, lambda, and R^2^ based on the full phylogenetic tree (n = 456). The best model is highlighted in bold with model comparison details outlined in Table 2.

|  | Estimate (95% BCI) | Posterior standard deviation |
| --- | --- | --- |
| Model 1: GSH ~ Activity; Lambda ($\lambda$) = 0.63, Bayesian $R^2$ value (95% BCI) = 0.67 (0.61–0.71) | | |
| Intercept | 0.03 (0.035 to 0.037) | 0.27 |
| Slope - Activity (CFAR) | 0.37 (0.25 to 0.48) | 0.06 |
| Model 2: GSH ~ Maximum body size; Lambda ($\lambda$) = 0.60, Bayesian $R^2$ value (95% BCI) = 0.65 (0.60–0.70) | | |
| Intercept | 0.028 (0.024 to 0.034) | 0.26 |
| Slope - Maximum body size | 0.32 (0.22 to 0.43) | 0.05 |
| Model 3: GSH ~ Median depth; Lambda ($\lambda$) = 0.69, Bayesian $R^2$ value (95% BCI) = 0.66 (0.61–0.71) | | |
| Intercept | 0.029 (0.024 to 0.037) | 0.3 |
| Slope - Median depth | -0.19 (-0.31 to -0.07) | 0.06 |
| **Model 4: GSH ~ Activity + Maximum size + Median depth; Lambda ($\lambda$) = 0.55, Bayesian $R^2$ value (95% BCI) = 0.66 (0.61 – 0.71)** |  |  |
| **Intercept** | **0.028 (0.024 to 0.034)** | **0.23** |
| **Slope - Activity (CFAR)** | **0.32 (0.21 to 0.43)** | **0.06** |
| **Slope - Maximum body size** | **0.23 (0.12 to 0.33)** | **0.05** |
| **Slope - Median depth** | **-0.19 (-0.3 to -0.08)** | **0.06** |

**Figure 2.**
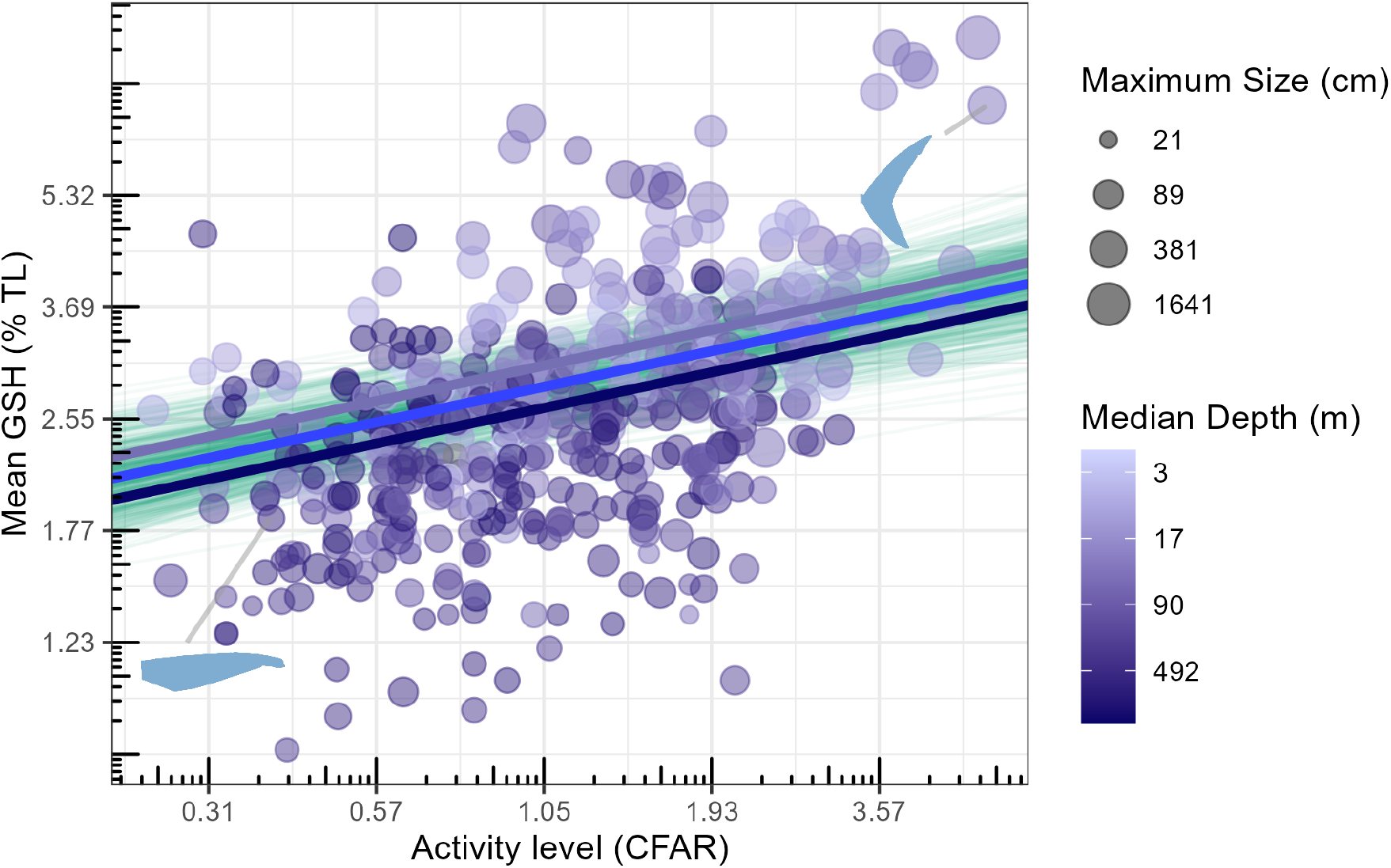
Relationship between activity level (Caudal Fin Aspect Ratio, CFAR) and mean gill slit height. Points are coloured by median depth and sized by maximum body size (Total length, cm). Green lines are 500 random draws of conditional fit from the posterior distribution of the global model with blue lines indicating intercept differences at varying depths; blue is mean depth (90 m), dark blue and light blue are 1SD in either direction (492 m and 17 m). Inset silhouettes are of the Longfin Mako Shark (*Isurus paucus*; top-right) and the Longfin Catshark (*Apristurus herklotsi*; bottom-left).

### Do larger-bodied sharks have larger gill slits?

GSH and maximum size were positively related such that species with larger maximum size had larger GSH. Maximum size ranged from the 15.7 cm Campeche Catshark (*Parmaturus campechiensis*) to the 2,000 cm Whale Shark (*Rhincodon typus*), with GSH values of 2.3% and 8.9% TL, respectively. An average shark species had a maximum size of 89 cm TL and an estimated GSH of 2.84 % TL (Table 1). GSH was estimated to increase by 0.33 sd (0.35% TL) for an increase in one sd of maximum size of (95 cm TL; Table 1).

### Do deeper-dwelling sharks have smaller gill slits?

GSH and median depth were negatively related such that deeper-dwelling species had smaller GSH (Figure 2 and S1). Median depth ranged from 2 m in the Northern Wobbegong (*Orectolobus wardi*) to 3,325 m in the Short-tail Catshark (*Parmaturus bigus*), with GSH values of 3.0% and 1.4% TL, respectively. An average shark with a median depth of 90 m would have an estimated GSH of 2.94% TL (Table 1). An increase in depth by one sd (402 m deeper) would result in a reduction in GSH of 0.19 sd (i.e., decrease by 0.2% TL; Table 1).

### Does ecological lifestyle explain GSH better than its components separately?

The effect of CFAR on GSH was positive and greater (β=0.31), than the effect of maximum size (β=0.23), whereas median depth had a negative effect on GSH (β= -0.17; Figure 2, Table 1). The global model had the most support out of the four models (*elpd_loo* = -458.25, Table 2). The models with the individual ecological lifestyle traits have less support than the global model, with the model with just CFAR being the second best model (Table 2). Nevertheless, the individual trait models estimated similar parameter estimates (grey coefficients; Figure 3) to the global model (black coefficients; Figure 3).

**Table 2.**
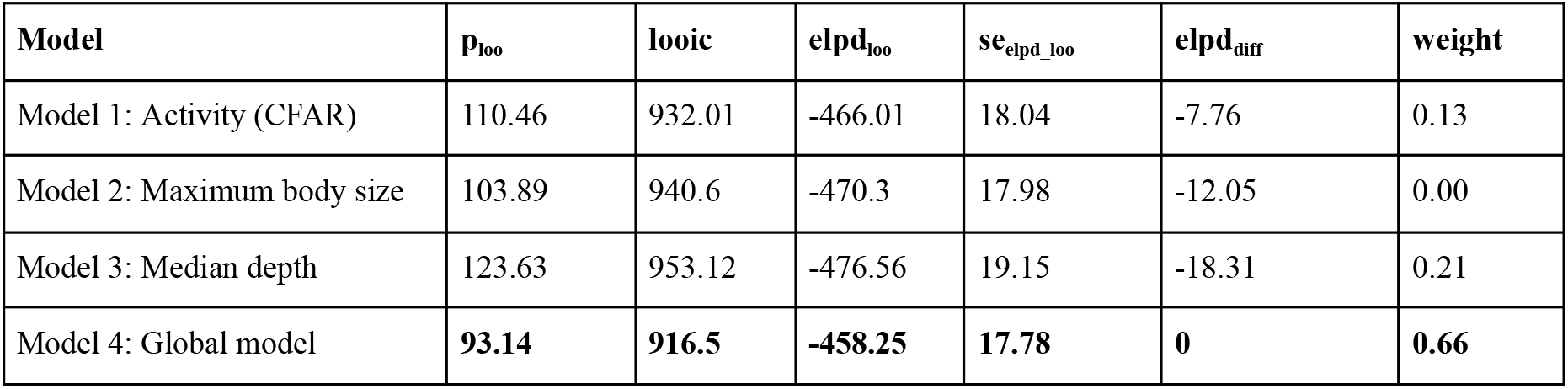
Model comparison table with the gill slit height models ordered as in Table 1, for full phylogenetic tree (n = 456)

| **Model** | **p<sub>loo</sub>** | **looic** | **elpd<sub>loo</sub>** | **se<sub>elpd_loo</sub>** | **elpd<sub>diff</sub>** | **weight** |
| --- | --- | --- | --- | --- | --- | --- |
| Model 1: Activity (CFAR) | 110.46 | 932.01 | -466.01 | 18.04 | -7.76 | 0.13 |
| Model 2: Maximum body size | 103.89 | 940.6 | -470.3 | 17.98 | -12.05 | 0.00 |
| Model 3: Median depth | 123.63 | 953.12 | -476.56 | 19.15 | -18.31 | 0.21 |
| Model 4: Global model | **93.14** | **916.5** | **-458.25** | **17.78** | **0** | **0.66** |

**Figure 3.**
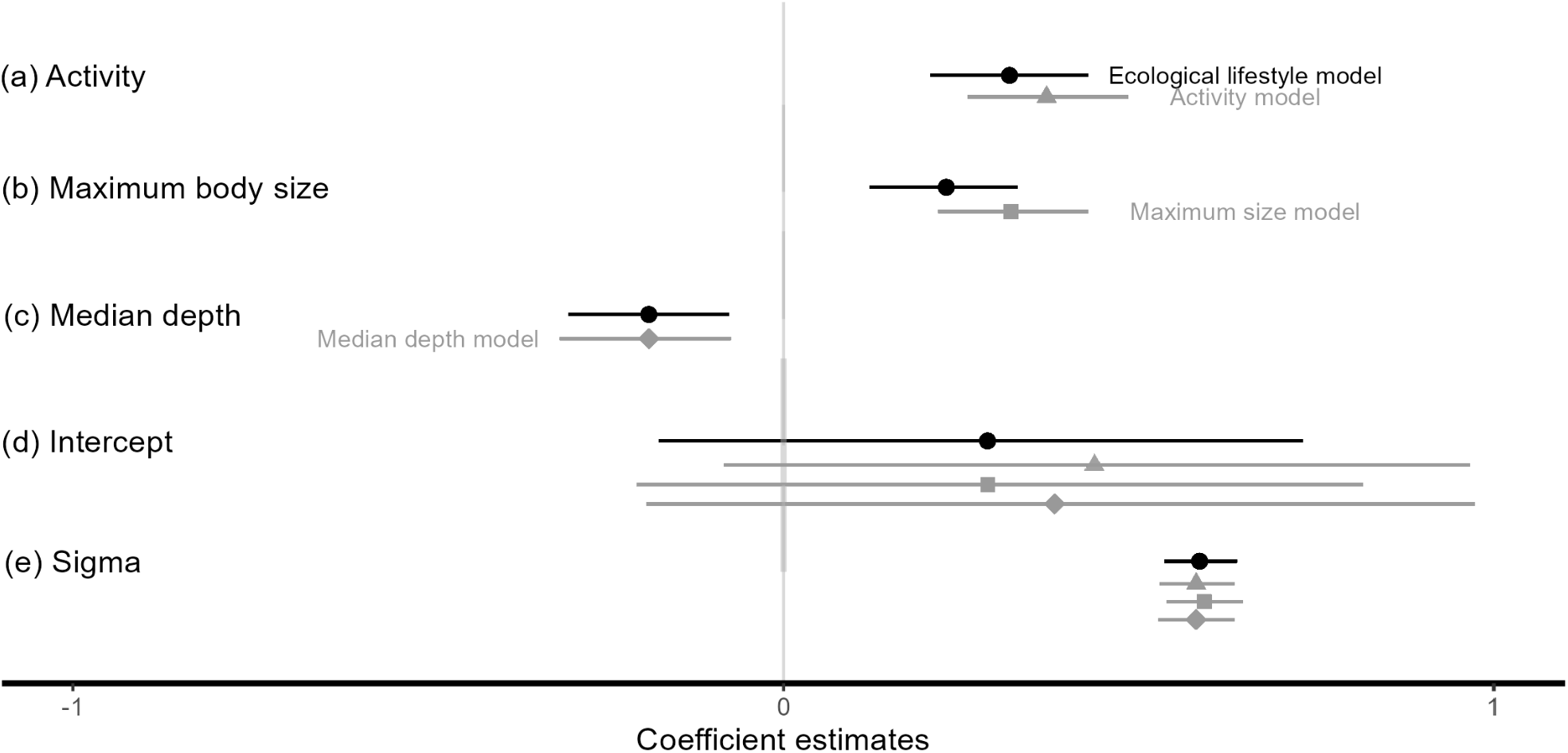
Model Coefficient estimates for each of the four models evaluated. Ecological lifestyle model estimates are in black (circles) while univariate model estimates are in grey (activity – triangles, maximum body size – squares, and median depth – diamonds). Lines for each coefficient indicate the 95% credible interval.

## 4 Discussion

Overall, we found that gill slit height was closely related to ecological lifestyle. Specifically, sharks with greater mean gill slit heights were more active (had greater caudal fin aspect ratios (CFAR), larger maximum sizes, and shallower depth ranges (Figure 2). Further, we found that the global model that considered all three ecological lifestyle traits together best explained gill slit height compared to any model with a given ecological lifestyle trait on its own. Next, we consider four points: (1) the connection between activity and oxygen demand, (2) the connection between maximum size and gill slit height, (3) the evolutionary response to reduced oxygen in the deep sea, and finally, (4) future directions.

### 4.1 Activity and oxygen demand

Activity level is long-known to be positively correlated with metabolic rate and gill surface area across fishes (Gray 1954, De Jager and Dekkers 1975, Bigman et al. 2018). Here, we show that activity level is also positively related to gill slit height across shark species (Figure 2), suggesting that we may only need to measure gill slits to infer activity and oxygen demand. One hypothesis that may explain these positive relationships is that species that are more active have a higher oxygen demand to support higher metabolic demand (Wegner et al. 2012, Killen et al. 2016, Bigman et al. 2023b).

### 4.2 Maximum body size and gill slit height

Larger-bodied species have larger gill slit heights and this relationship hints at a more general one between gill morphology and body size across sharks. GSH in this study is scaled to be proportional to total length, so sharks with greater GSH have larger GSH in relation to their body length. In this sense, if each species had the same total length, variation in GSH may correspond to differences in maximum or asymptotic sizes, with larger-bodied species having larger GSH and smaller-bodied species having smaller GSH. This may indicate that species with larger GSH are expending extra resources on maintaining larger gills to potentially reach larger asymptotic sizes (i.e. facilitate growth). The ontogenetic scaling of gill surface area within each shark species is about 0.85 (range 0.75–0.9), regardless of their maximum size (Bigman et al. 2018, 2023a, b). Further, species that grow to larger maximum sizes maintain larger than average gills throughout their lifetime, which allows for the necessary oxygen uptake for growth, maintenance and reproduction (Bigman et al. 2018).

### 4.3 The evolutionary response to reduced oxygen in the deep sea

Deepwater sharks could provide some insight into the connection between metabolic rate and functional morphology. Species that have evolved in deepwater have made tradeoffs between metabolic processes to ensure their demands do not exceed the environmental conditions. Deepwater species tend to be smaller-bodied with smaller gill slit heights and low activity levels (lower CFAR; Figure 2 darker shades of blue). There are multiple abiotic challenges to living in the deepsea (>200 m), most notably the low temperature and oxygen concentration. Species that live in deeper waters are often in environments with much lower temperatures and oxygen levels than pelagic and coastal surface waters (Garcia et al. 2008; Rigby & Simpfendorfer 2015; Pardo & Dulvy 2022).

The evolutionary response to reduced oxygen in the deep sea could be (i) increased gill area, (ii) reduced body size (hence growth rates and reproduction), or (iii) reduced activity, or some combination of the three responses. Generally, in contrast to the first idea above, deepwater sharks have shorter gill slit heights. However, in some species, gill slit height is elongated by extending the gill slits under the body to end only at the midline, as seen in Frilled Sharks (Family Chlamydoselachidae) and the filter-feeding Basking Shark (*Cetorhinus maximus*). A small number of deepwater shark species have increased the number of parabranchii and gill slits, as seen in the Cow Sharks (Hexanchidae), such as the Bluntnose Sixgill Shark (*Hexanchus griseus*), and Broadnose Sevengill Shark (*Notorhynchus cepedianus*), as well as the Sixgill Stingray (*Hexatrygon bickelli*). While these species have additional gill slits using their median gill slit height compared to accounting to summed gill slit height has little effect on our overall results (Figure S3, S4; Table S4). Another way to increase gill area is to increase head size laterally to accommodate larger gill surface area is exemplified in the Lollipop Shark (*Cephalurus cephalurus*) and Filetail Catshark (*Parmaturus xaniurus*), which inhabits anoxic water (with oxygen concentrations of <1% of that found in surface waters; 1.6 μmol/kg), and as such, is a low-oxygen specialist (Ebert et al. 2021). Similarly, the Bigeye Thresher (*Alopias superciliosus*) is the only thresher that forages in the deep Oxygen Minimum Zone and has the largest known gill surface area of any elasmobranch with an expanded branchial cavity to accommodate the larger gill surface area (Wootton et al. 2015). So while some species have an extra gill arch or two or enlarged heads to cope with anoxia, overall, the general pattern is of smaller, rather than larger gill slit heights (and presumably gill surface area), in deeper sharks.

Second, additional adaptations to life in low oxygen environments include reducing metabolic demand by reducing size or activity. Reduced metabolic rate of deepwater species is thought to underlie reduced growth rates and reproductive output compared to shallow-water species (Drazen & Seibel 2007, Pardo & Dulvy 2022). Deepwater sharks tend to be small, slow growing, have low reproductive output, and generally, have slower life histories (Simpfendorfer & Kyne 2009, Pardo & Dulvy 2021). Deepwater habitats exhibit some of the smallest shark species, suggesting that over evolutionary time, deepwater species have reduced their maximum size to ensure they are not in a constant state of hypoxia in deeper water. Third, deepwater sharks tend to have much lower activity levels and swimming speeds (Pinte et al. 2020). Indeed, the most active deepwater sharks, the lantern sharks (Etmopteridae), are also among the smallest species of sharks in the world (Pinte et al. 2020), suggesting deepwater sharks can be small and active or large and inactive, but not large-bodied and active.

Another possible limiting factor in the deep is prey availability and encounter rates (visual-interactions hypothesis, Drazen & Seibel 2007). With reduced (or no) visible light, the chances of encountering prey is much lower than in shallow, light-filled waters and, thus, reduces resources for metabolism. The reduced resources at depth and energetic costs of a lipid-rich and buoyant liver in chondrichthyans may explain why they are largely absent below 3000 m (Treberg & Speers-Roesch 2016). Together, the low temperatures, low oxygen, and low food availability could explain the smaller gills and slower pace of life observed in deep sea sharks as well as innovations such as the evolution of more gill slits and larger heads (Pardo & Dulvy 2022). Clearly, understanding the evolutionary trajectory of these adaptations is a key question, and according to the Compagno (1990) hypothesis, deepwater sharks are ancestral and hence the hypothesis to test is whether the low activity, smaller gill lifestyle is ancestral and the general evolutionary trajectory has been toward more active larger gill lifestyles of coastal and pelagic species.

### 4.4 Future directions

Although gill slit heights measured from anatomically-accurate illustrations in field guides are comparable to those measured on physical specimens, we recognize that our morphological measurements (GSH and CFAR) from such illustrations may not be identical to those from physical specimens. We have confirmed the relationship between gill slit height of field guide drawings closely matches laboratory measurements in five species of carcharhinid (Vanderwright et al. 2020). Field guides are routinely used as a source of data in scholarly publications, and specifically, ecological studies (Schmidt 2006). Such approaches offer the advantage of requiring relatively low effort and cost, and can be applied consistently and repeatedly across the whole lineage. There are three alternative validation approaches that are expensive and laborious, but nevertheless complementary and would build trust in these simple trait measures. First, by opportunistically measuring live specimens or individuals that have been recently captured from ongoing fieldwork, possibly sourced through a ‘citizen’ science project. Second, museum specimens can be used to validate the accuracy of field guide illustrations but they have been preserved for many years and may have experienced shrinkage from fixing agents as well as damage from multiple specimens in cramped storage containers. Third, there is increasing development of underwater laser photogrammetry measuring techniques, to estimate maximum size using paired lasers to provide a reference size within the field-of-view of a video camera. These data could also be used to measure gill slit height and CFAR (Marshall et al. 2011, Meekan et al. 2006).

### 4.5 Conclusion

We have developed simple, easily measured, cost-effective trait measures relationships linking metabolic morphology and ecological lifestyle across sharks. These measures can help shed light on the metabolic ecology of a large lineage of marine fauna that have very few and hard to gather metabolic and life history data (Lyons et al. 2019). Our previous work has shown that metabolic rates are weakly related to life history traits, such as growth rate and age at maturation, however the exploration of these relationships is constrained by the limited availability of metabolic rate data (Wong et al. 2021; Gravel et al. *In Review*). We hope that these metabolic morphology traits will improve predictions of life history traits and maximum intrinsic population growth rate, while minimizing the collection of live specimens or spending countless hours in laboratories. Many species are data-poor, particularly on the population scale at which management occurs (Kindsvater et al. 2019). The development of metabolic morphology traits will go a long way to expand the breadth of species that can be evaluated using Ecological Risk Assessments including species that are rarely observed or have only been observed a small number of times.

## Acknowledgements

We thank Will Stein and Chris Mull for helping with phylogenetic trees. We also thank Sarah Gravel, Amanda Arnold, Rachel Aitchison, Jay Matsushiba, Brogan Neufeld, Hannah Watkins, and the anonymous reviewers for their constructive comments on this manuscript.

