## Supplementary materials for "Ecological lifestyle and gill slit height across sharks"

### Supplementary Figures

**Figure S1.** Relationship of median depth and mean gill slit height across sharks. Dashed lines indicate trait means in the dataset, so the horizontal line indicates the average mean gill slit height is 3% of the maximum body length and the average median depth is 90 metres.

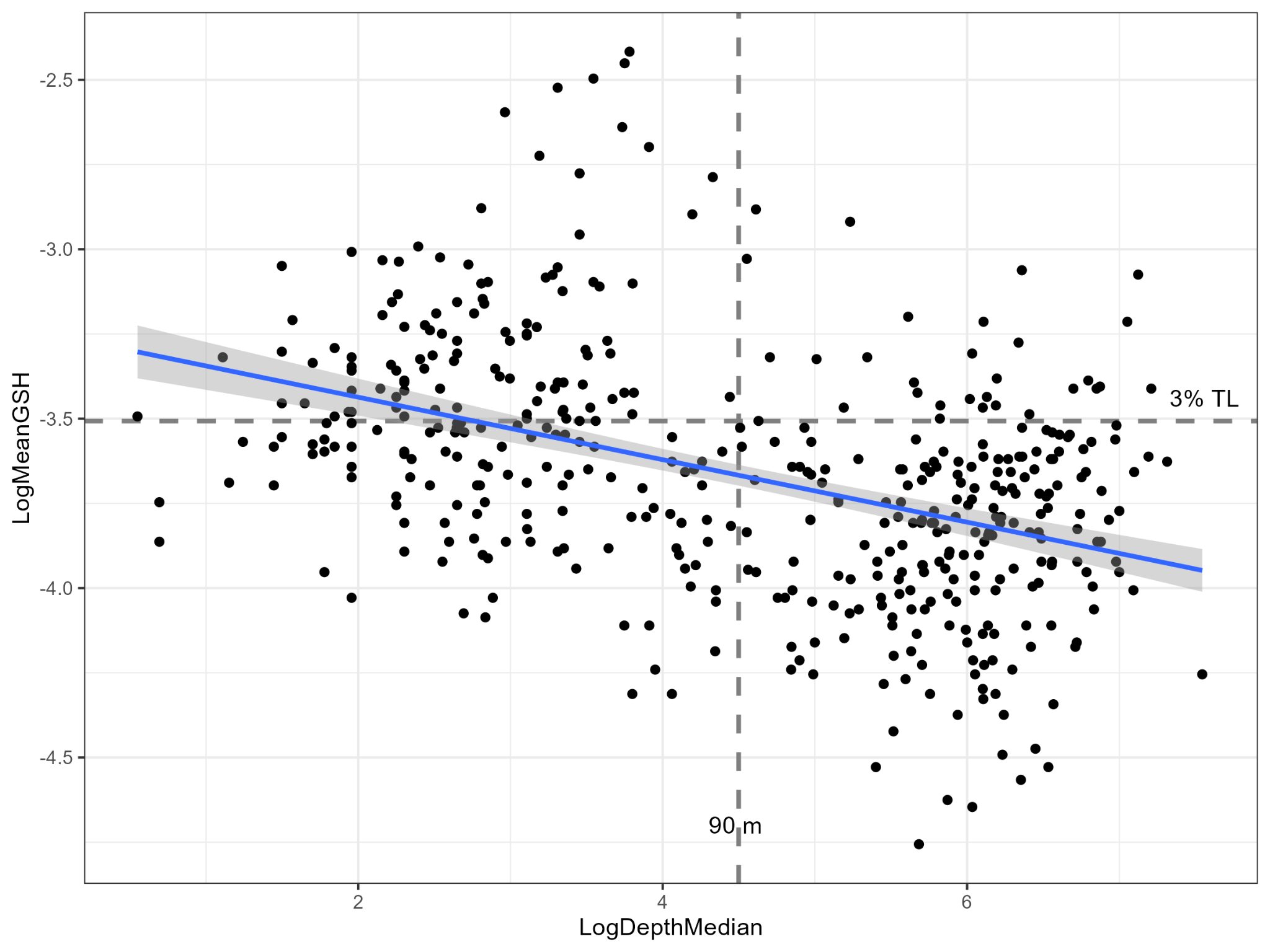

**Figure S2.** The negative relationship between median depth and maximum body size.Dashed lines indicate trait means in the dataset, so the horizontal line indicates the average maximum body size is 89 cm TL and the average median depth is 90 metres.
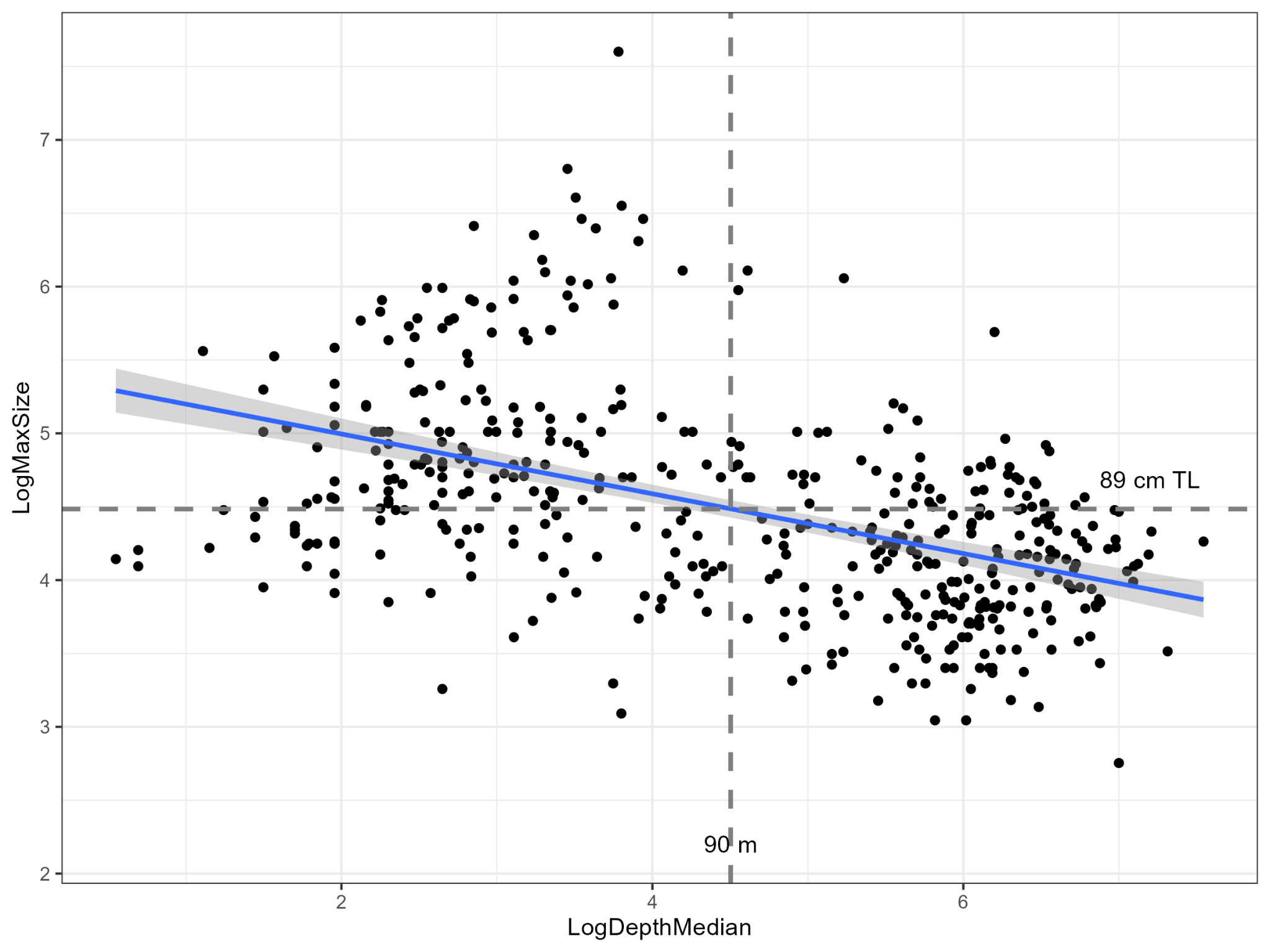

**Figure S3.** Relationships are insensitive to the choice of mean gill slit height (left hand column) and summed gill slit height (right hand column, versus activity (CFAR, upper row), maximum body size (middle row), and median depth (lower row)

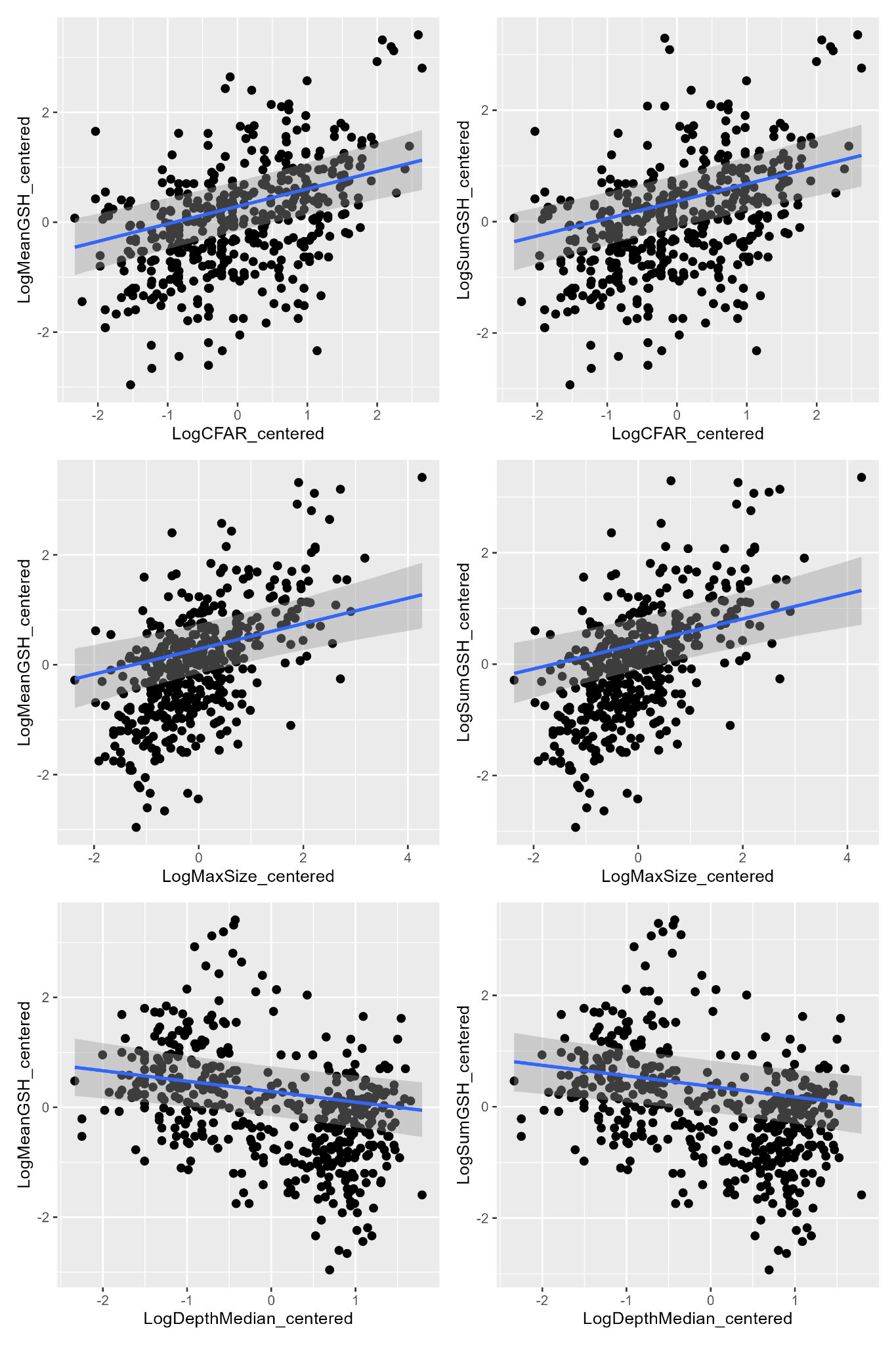

**Figure S4.** Relationship between gill slit height and activity (CFAR) which the difference in standardized mean gill slit height (grey dots) and the standardized summed gill slit height for the cow sharks (Hexanchidae), i.e. the 6- and 7-gilled sharks. The three thresher sharks are illustrated in blue.

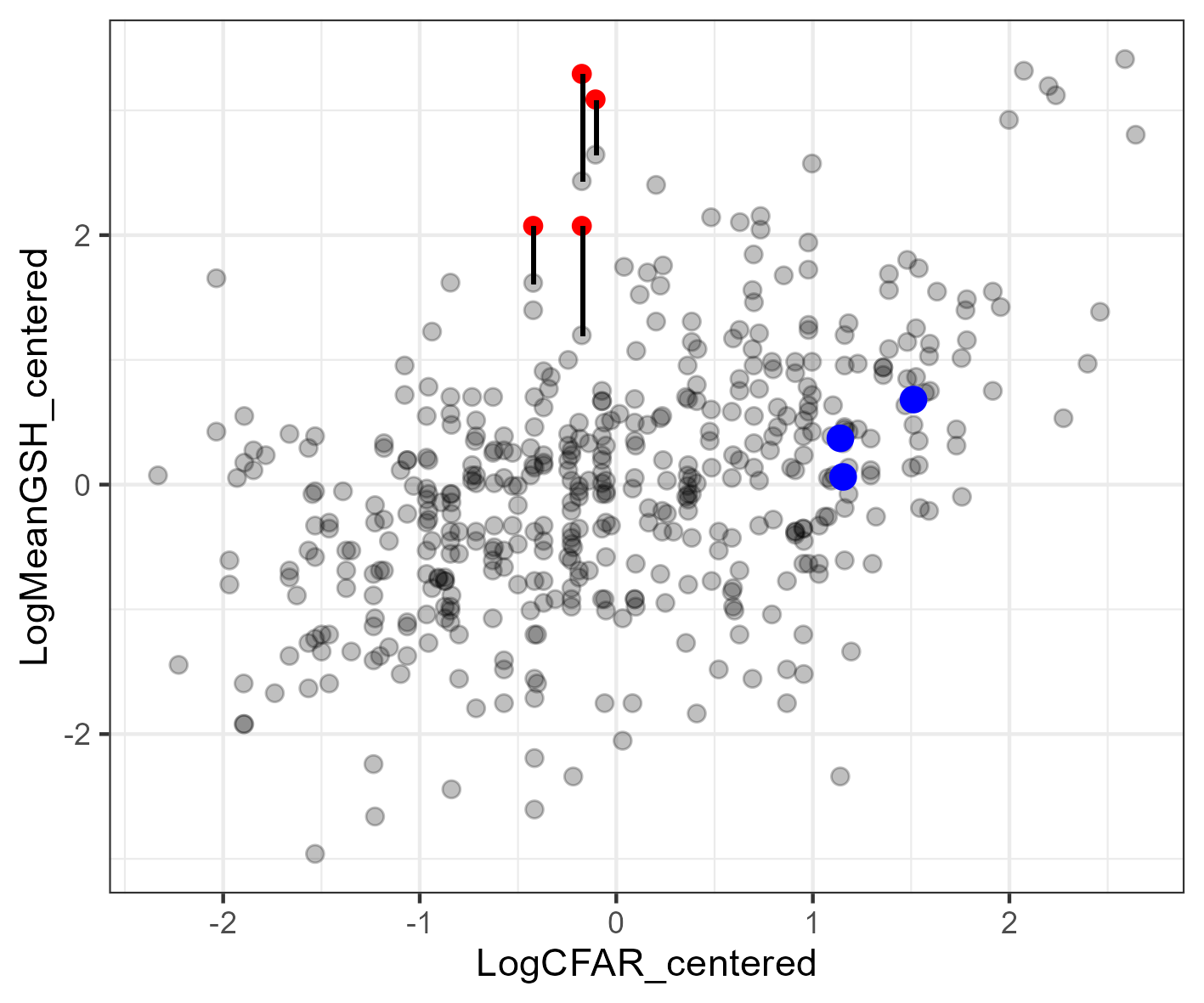

### Supplementary Tables

**Table S1.** Model summary table comparing parameter estimates (with bayesian credible intervals), posterior standard deviation, lambda, and R^2^ based on the molecular phylogenetic tree (n = 268).

|  | Estimate (95% BCI) | Posterior standard deviation |
| --- | --- | --- |
| Model 1: GSH ~ Activity; Lambda (𝜆) = 0.69, Bayesian R^2^ value (95% BCI) = 0.73 (0.66 – 0.79) | | |
| Intercept | 0.034 (0.028 to 0.042) | 0.28 |
| Slope - Activity (CFAR) | 0.37 (0.22 to 0.51) | 0.07 |
| Model 2: GSH ~ Maximum body size; Lambda (𝜆) = 0.65, Bayesian R^2^ value (95% BCI) = 0.72 (0.65 – 0.78) | | |
| Intercept | 0.032 (0.026 to 0.039) | 0.27 |
| Slope - Maximum body size | 0.38 (0.24 to 0.51) | 0.07 |
| Model 3: GSH ~ Median depth; Lambda (𝜆) = 0.74, Bayesian R^2^ value (95% BCI) = 0.73 (0.66 – 0.80) | | |
| Intercept | 0.032 (0.025 to 0.041) | 0.32 |
| Slope - Median depth | -0.16 (-0.32 to 0.01) | 0.08 |
| Model 4: GSH ~ Activity + Maximum size + Median depth; Lambda (𝜆) = 0.61, Bayesian R^2^ value (95% BCI) = 0.72 (0.65 – 0.78) | | |
| Intercept | 0.032 (0.027 to 0.039) | 0.25 |
| Slope - Activity (CFAR) | 0.29 (0.15 to 0.43) | 0.07 |
| Slope - Maximum body size | 0.29 (0.16 to 0.42) | 0.07 |
| Slope - Median depth | -0.17 (-0.32 to -0.02) | 0.08 |

**Table S2**. Variance Inflation Factor table for the global model without random effect of the phylogenetic tree

| **Parameter** | **VIF** |
| --- | --- |
| Activity (CFAR) | 1.39 |
| Maximum body size | 1.72 |
| Median depth | 1.34 |

**Table S3.** Correlation table of model parameters examined in this study

|  | **Mean GSH** | **CFAR** | **Maximum body size** | **Median depth** |
| --- | --- | --- | --- | --- |
| **Mean GSH** | 1.00 |  |  |  |
| **CFAR** | 0.47 | 1.00 |  |  |
| **Maximum body size** | 0.61 | 0.53 | 1.00 |  |
| **Median depth** | -0.45 | -0.27 | -0.50 | 1.00 |

**Table S4.** Model estimates for the global model fitted with summed GSH as the response variable (i.e., Summed GSH ~ Activity + Maximum body size + Median Depth). See Figure S3 for visual comparison.

|  | **Estimate** | **Estimate Error** |
| --- | --- | --- |
| **Intercept** | 0.036 (-0.01 to 0.08) | 0.24 |
| **Activity (CFAR)** | 0.31 (0.20 - 0.42) | 0.06 |
| **Maximum body size** | 0.22 (0.12 to 0.33) | 0.05 |
| **Median depth** | -0.19 (-0.30 to -0.08) | 0.06 |
